# EpiFusion: Joint inference of the effective reproduction number by integrating phylodynamic and epidemiological modelling with particle filtering

**DOI:** 10.1101/2023.12.18.572106

**Authors:** Ciara Judge, Timothy Vaughan, Timothy Russell, Sam Abbott, Louis du Plessis, Tanja Stadler, Oliver Brady, Sarah Hill

## Abstract

Accurately estimating the effective reproduction number (Rt) of a circulating pathogen is a fundamental challenge in the study of infectious disease. The fields of epidemiology and pathogen phylodynamics both share this goal, but to date, methodologies and data employed by each remain largely distinct. Here we present EpiFusion: a joint approach that can be used to harness the complementary strengths of each field to improve estimation of outbreak dynamics for large and poorly sampled epidemics, such as arboviral or respiratory outbreaks, and validate it for retrospective analysis. We propose a model of Rt that estimates outbreak trajectories conditional upon both phylodynamic (time-scaled trees estimated from genetic sequences) and epidemiological (case incidence) data. We simulate stochastic outbreak trajectories that are weighted according to epidemiological and phylodynamic observation models and fit using particle Markov Chain Monte Carlo. To assess performance, we test EpiFusion on simulated outbreaks in which transmission and/or surveillance rapidly changes and find that using EpiFusion to combine epidemiological and phylodynamic data maintains accuracy and increases certainty in trajectory and Rt estimates, compared to when each data type is used alone. Finally, we benchmark EpiFusion’s performance against existing methods to estimate Rt and demonstrate advances in efficiency and accuracy. Importantly, our approach scales efficiently with dataset size, including the use of phylogenetic trees generated from large genomic datasets. EpiFusion is designed to accommodate future extensions that will improve its utility, such as introduction of population structure, accommodations for phylogenetic uncertainty, and the ability to weight the contributions of genomic or case incidence to the inference.

**Author Summary:** Understanding infectious disease spread is fundamental to protecting public health, but can be challenging as disease spread is a phenomenon that cannot be directly observed. So, epidemiologists use data in conjunction with mathematical models to estimate disease dynamics. Often, combinations of different models and data can be used to answer the same questions – for example ‘traditional’ epidemiology commonly uses case incidence data (the number of people who have tested positive for a disease at a certain time) whereas phylodynamic models use pathogen genomic sequence data and our knowledge of their evolution to model disease population dynamics. Each of these approaches have strengths and limitations, and data of each type can be sparse or biased, particularly in rapidly developing outbreaks or lower-middle income countries. An increasing number of approaches attempt to fix this problem by incorporating diverse concepts and data types together in their models. We aim to contribute to this movement by introducing EpiFusion, a modelling framework that makes improvements on efficiency and temporal resolution. EpiFusion uses particle filtering to simulate epidemic trajectories over time and weight their likelihood according to both case incidence data and a phylogenetic tree using separate observation models, resulting in the inference of trajectories in agreement with both sets of data. Improvements in our ability to accurately and confidently model pathogen spread help us to respond to infectious disease outbreaks and improve public health.

## 1. Introduction

The effective reproduction number (Rt) is a helpful epidemiological parameter for characterising disease transmission. Rt refers to the time-varying average number of secondary infections resulting from a primary infected individual and can vary due to factors such as population immunity, human behaviour, or changes in pathogen infectiousness. Retrospective modelling of how Rt varies over the course of an outbreak, such as in the examples we outline in this paper, allows for evaluation of policy and intervention efficacy (1–4), and quantifying how different factors contribute to Rt can inform outbreak preparedness planning by providing the basis for modelling spread under different scenarios (5). Classical epidemiology (6) and phylodynamics (4) often aim to infer Rt but use distinct methodologies and data to achieve this goal. Both fields face similar but non-overlapping obstacles in terms of data availability, reliability, and bias (7–10). We investigate an approach to estimate Rt that reduces this uncertainty through linking principles of phylodynamic and epidemiological modelling using particle Markov Chain Monte Carlo (pMCMC) (11) which is scalable to large datasets.

Phylodynamic approaches allow estimation of the genealogical history of sequenced virus samples and can therefore inform about disease spread that occurred prior to the first identified case. Phylogenetic trees frequently capture unusual population dynamics (12) that are not normally detectable using case data alone, such as long-range virus lineage movements, or importations or growth in specific variants. However, a central challenge for phylodynamics is that genomic data sampling density can be low or spatiotemporally biased relative to infection occurrence (13). Furthermore, Rt has thus far been commonly estimated as a piecewise constant function that rarely has sufficient temporal resolution to be useful for public health decision making (14), with some exceptions (15).

Conversely, epidemiological models of Rt use case data that are often more temporally consistent than genomic data, and usually have greater flexibility than phylodynamic models to accommodate additional information such as climatic or human movement data (16–19). However, case data can be easily biased by changes in case definitions or reporting practices (8,20), which can cause artificial fluctuations in Rt estimates, and there is no data to examine disease dynamics prior to the first case observation (whereas phylogenetic tree data can be used to reconstruct past pathogen dynamics). Furthermore, viruses that can cause similar clinical symptoms (such as Zika, chikungunya and dengue viruses (21,22)) can be easily misreported where specific molecular or serological testing is not conducted. This can result in the inferred Rt capturing the average population dynamics of multiple cocirculating pathogens, which is less useful to inform disease-specific control measures such as vaccination programs (23–25).

Notably, phylodynamic and epidemiological approaches may vary in their effectiveness at different stages of an outbreak (13). Approaches that combine principles and data from both phylodynamic and epidemiological models could improve our ability to estimate Rt, by taking advantage of the complementary strengths of each field.

Early attempts to use both phylodynamics and epidemiology to estimate disease dynamics typically employed a ‘corroborate or contradict’ strategy, where methods and data native to each field were used separately to address the same research question (26–28). Alternatively, methods from each field have sometimes been used to address different research questions in the same study (29). Recently, attempts have been made to progress joint inference approaches that use both phylodynamic (dated genomic sequence) and epidemiological (case incidence) data as input to a single model (30–33). Many of these attempts have built on the principle of the particle filter (11). Particle filtering is a sequential Monte Carlo approach that aims to approximate the posterior distribution of a state variable in a stochastic process (in this case, an epidemic trajectory). Particles move through a hidden Markov Model (the process model) and are weighted by their likelihood according to the data (the observation model). They can then be resampled according to their weights, resulting in the propagation of particles with estimated states consistent with the data under an observation model. The use of particle filtering is arguably the most straight-forward method to directly link epidemiological and phylodynamic models, as the resampling of particles through time allows the genealogical and epidemiological data to jointly influence the particle states during the state-simulation process.

Particle filtering is well established for use with epidemiological case incidence data, and there are many existing implementations of particle filtering in epidemiological modelling (34,35). More recently, appropriate particle filtering approaches have been developed that can use genealogies obtained from sequence data. Rasmussen et. al first proposed a joint inference approach consisting of a common process model and separate observation models for a genealogy and case incidence data (31). This methodology was later extended to allow fitting of epidemiological models that incorporate simple population structure (32), and was also used as the basis of an approach for inferring transmission heterogeneity (36). These models were all reliant on coalescent-based phylodynamic methods and assumed independence between case incidence and events in the phylogenetic tree (37). In 2019 Vaughan et. al proposed a method ‘EpiInf’, that enable use of birth-death phylodynamic methods within a particle filter to infer epidemic trajectories through time (33). EpiInf derived a phylodynamic likelihood that explicitly models case incidence data as ‘unsequenced observations’ within the phylodynamic observation model as ‘events’ on the tree, thus overcoming the independence assumption made in earlier approaches. However, this latter approach does not scale well, as it quickly becomes intractable as the number of sequences or cases increases (even when using a tau-leaping approximation (38)). It also greatly limits the possible complexity that could be obtained using a separate epidemiological observation model, which could feasibly incorporate diverse data sources (e.g. climate or human movement data). Conversely TimTam (30), proposed by Zarebski et al. in 2022, is a (non-particle filtering) birth-death phylogenetic approach that can integrate case incidence and genomic sequence data in a computationally tractable way by approximation of the birth-death observation model density (39,40), while also eliminating the assumption of independence between tree and occurrence events. However, while it is possible to infer prevalence at user specified times and Rt in piecewise constant intervals, it is not possible to infer continuous epidemic trajectories with this model, which limits its ability to detect transmission fluctuations at a higher temporal resolution.

We investigate a new approach, EpiFusion, that extends existing implementations that employ particle filtering or pMCMC (31,33) to reconstruct epidemic trajectories using case incidence and a phylogenetic tree either individually or together, while making the assumption that the tree and case trajectory are independent. Our proposed approach improves on limitations of previous methods by *(i)* introducing a birth-death based phylodynamic likelihood to a dual observation model structure *(ii)* making improvements in computational efficiency and *(iii)* allowing epidemic trajectories to be inferred in greater temporal resolution.

## 2. Methods

We adopt an overall structure based on the ‘common process model – dual observation model’ structure *(Figure 1)* used by Rasmussen et. al (31) and validated by many particle filtering methods outside of the context of infectious disease (41,42). The data inputs to this model are case incidence, a time-scaled phylogenetic tree constructed from virus genomic sequences, or both datasets together. In practice the frequency of the case incidence data could be user specified, (daily, weekly, monthly etc.), but for the examples shown here we use a weekly resolution. The performance of the model using these different data sources independently and their combination is compared during model validation.

**Figure 1.**
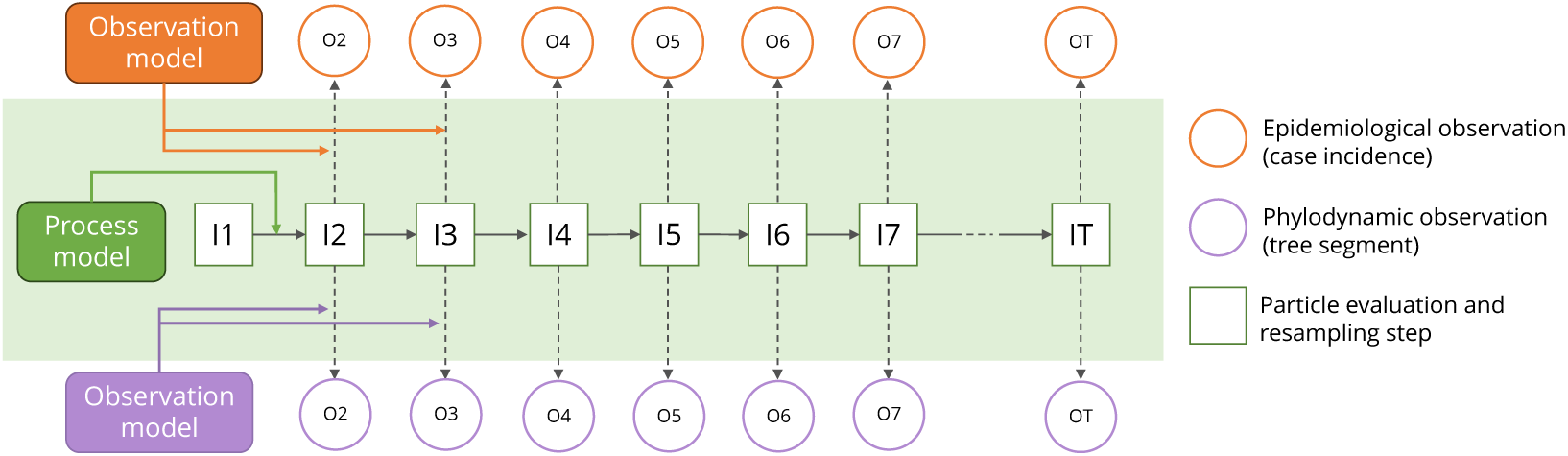
Overall EpiFusion particle filter structure, with the particle states (green outlined boxes) driven by the parameters of the process model, and evaluated at resampling steps by epidemiological and phylodynamic observation models against case incidence and phylogenetic tree segments respectively (orange and purple circles).

### Process Model

We use the term ‘process model’ to define how particle states are incremented between resampling steps in the particle filter. Particles model the number of infected individuals (*I*) in discrete daily intervals driven by a process model assuming independent Poisson-distributed infection and recovery counts (*Eqn 1*).

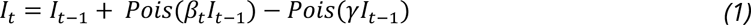

Previous particle filtering approaches have modelled more complex population compartment structures such as SIR or SIS for the particle states, and simulated trajectories in continuous time (i.e. simulating each infection and recovery event individually). We have implemented this daily discretisation and solely modelled the ‘I’ compartment as a trade-off that allows improved computational efficiency. Transmission dynamics that would be driven by the S and R compartments are instead captured by varying the infection rate *β* over time (see ‘Fitting with MCMC’).

There are two methods for deriving Rt trajectories from this process. The first is by storing new daily infections (*Pois*(*β*_t_*I*_t−1_)) and using a renewal equation (43) with a user-defined probability mass function of the pathogen generation time. Alternatively, Rt can be calculated using the values of *β* and *γ*, i.e. 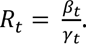 For this paper we use the former approach to facilitate direct comparison to the simulated dataset by performing the renewal equation on the simulated true infection trajectory to obtain the ‘true Rt’.

### Observation Models

At each resampling step, the particle states are evaluated against epidemiological and phylogenetic data using individual ‘observation models’; that is, the models that define the weights (*ω*) of each particle state according to each dataset.

The provided epidemiological data, here weekly reported case counts (*ct*), are evaluated using a Poisson probability density function and the epidemiological sampling rate *φ* to give the ‘epidemiological weight’ of the particle *(Eqn 2)*.

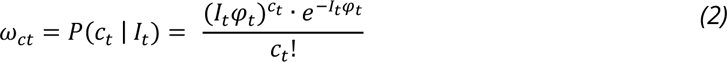

The particle weight given the phylodynamic data (a one-day segment of a time-scaled phylogenetic tree *gt*) is a daily discretisation of that which was derived by Vaughan et. al for EpiInf (*Eqn 3*). We implement an importance sampling strategy to prevent trajectory events that are incompatible with the tree structure (for example recovery events that result in fewer individuals being infected than there are lineages in the tree). Thus, the phylodynamic weight is the product of the conditional probability of the tree given the compartment size *P*(*g*_t_|*I*_t_) with the ratio of the probability of the trajectory under the process model, *P*(*I*), to the probability of the trajectory under the importance distribution *P*′(*I*). This is the sum of the probabilities of the observed events (number of observed infections of new individuals *bt*; number of sampling events *st*) on the tree, the exponentially distributed waiting times for events that were not observed (infections with rate *β*^°^t*I*_t-1_ and genomic samplings of infected individuals with rate *ψI*_t-1_), and the forbidden recovery propensity *γ*^f^t*I*_t-1_. Further information on the importance sampling implementation is available in the Supplementary Information (page 37).

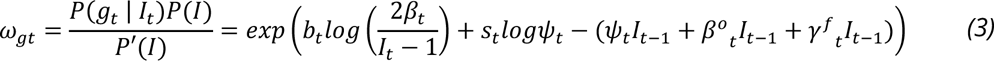

During resampling, the particles are weighted (*ω*_particle_) by the product of their phylodynamic and epidemiological weights (*Eqn 4*), thus facilitating the propagation of particles that are consistent with both the phylogenetic and epidemiological data.

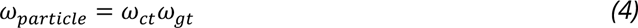

### Fitting with MCMC

Following completion of the particle filter, the overall likelihood of each estimated trajectory across the whole outbreak consists of the product of the average particle weights at each resampling step (*Eqn 5*). This is therefore the likelihood of a particle trajectory under the importance distribution sampled from the surviving particles given the epidemiological and phylogenetic data, and the parameter set of the particle filter *θ* which can be fit using MCMC.

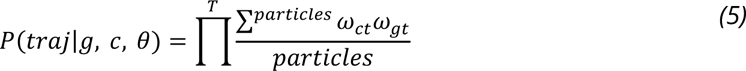

This model is fit using Metropolis-Hastings MCMC sampling, deriving posterior samples of the number of infected individuals over time, and the rates *β*, *γ*, *φ* and *ψ*. For more complex analyses where these rates change over time, the parameters fit by the MCMC estimate these timewise rates. Three options are available for defining and fitting a time-varying rate; (i) the rate at time *t* is a function of a logistic curve, and the curve parameters are fit via MCMC, (ii) the rate is defined as a piecewise constant function for which the value in each interval is fit via MCMC, and for which interval times can be either pre-specified or also inferred via MCMC, and (iii) the rate is treated as part of the particle state and fit in the particle filter as a random walk, with the initial rate value and standard deviation of the random walk fit via the MCMC. For the examples shown in this paper, the latter approach (iii) was used.

### Model Validation

To validate the model we simulated various outbreak scenarios (*Table 1*) in a population of 20,000 individuals under different conditions using ReMASTER (44,45), and specifically highlight three of these scenarios here. Parameters to simulate realistic outbreaks were chosen based on literature documenting various large, poorly sampled outbreaks (46–55). The simulations can be divided into the introduction of a novel pathogen into an immune naïve population with constant sampling, an introduction scenario with a step-change in sampling when the outbreak is ‘discovered’, and a step change in transmission of an endemic pathogen. The simulated datasets also explored different population compartment compositions: SEIR, SIR, and SIRS respectively. For scenarios *(i)* and *(ii)*, the scenario was initiated with a fully susceptible population and one infected individual, but scenario *(iii)* was initiated with a lower number of susceptible individuals to quickly reach an equilibrium that simulated endemic transmission. Simulated weekly reported case incidence counts and phylogenetic trees of samples were produced from each scenario (*Figure 2*), and the true prevalence according to the simulation was known in continuous time. The model was evaluated based on its ability to reconstruct from the simulated data the true infection incidence and effective reproduction number over time, as well as other epidemiological parameters (e.g. *γ*, *φ*, *ψ*) of the scenario, from the simulated data. To assess the advantage of combining phylodynamic and epidemiological data in this framework, models using solely the phylogenetic tree or case incidence data were compared to the combination of both. The models were also benchmarked against existing packages EpiNow2 (56), BDSky (14) and TimTam (30,57) with the same simulated data. For BDSky and TimTam, which take a sequence alignment as their input rather than a phylogenetic tree, sequences were simulated from the ReMASTER tree using the R package phangorn (58).

**Figure 2.**
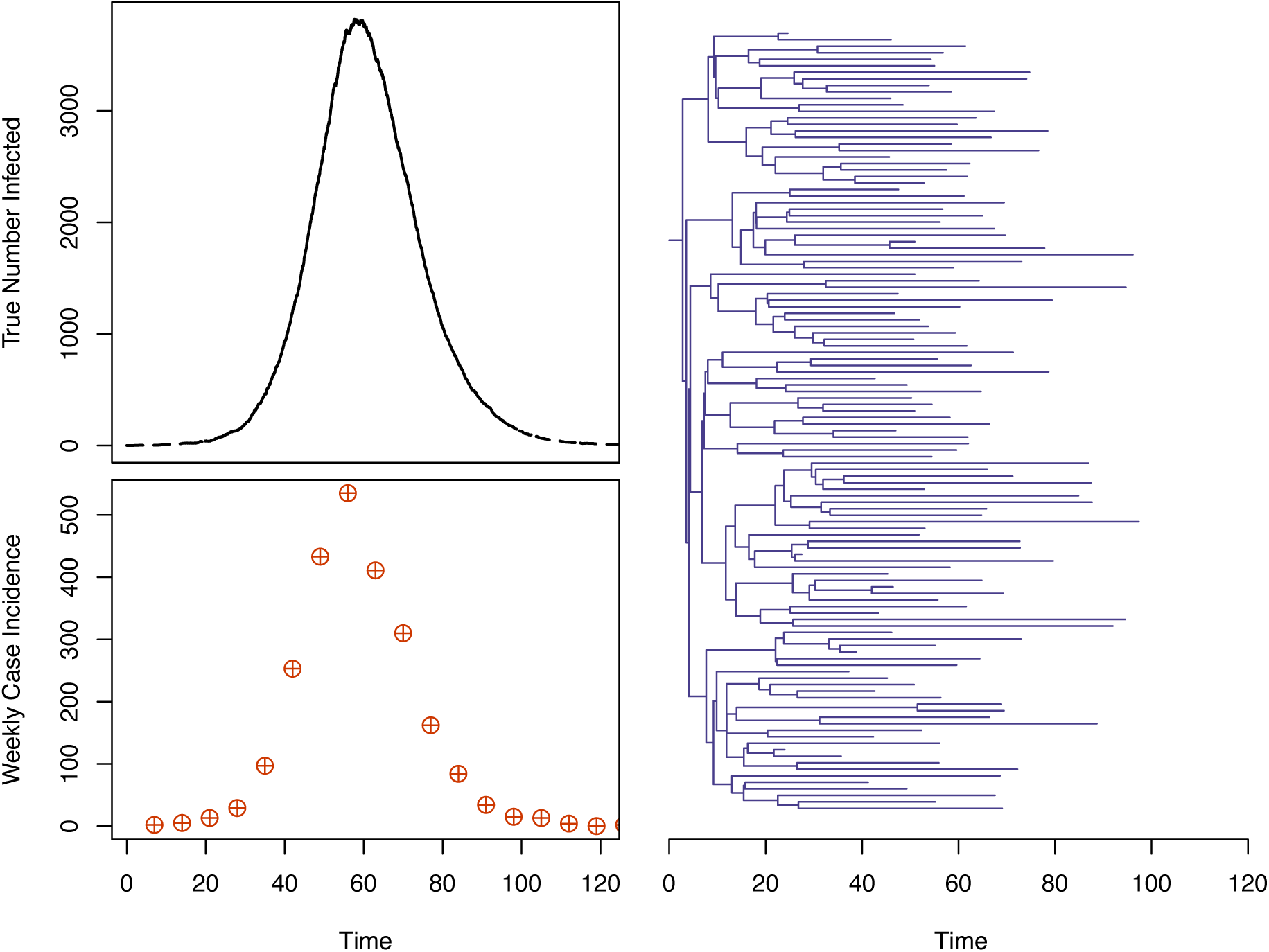
Simulated epidemic (introduction scenario) for model validation, and corresponding data for fitting. True number of people infected (top left) demonstrates the epidemic peak, from which weekly reported case incidence counts (bottom left) and a phylogenetic tree (right) of simulated samples were derived. Plots of the other simulated datasets are included in the Supplementary Information (page 39).

**Table 1.**
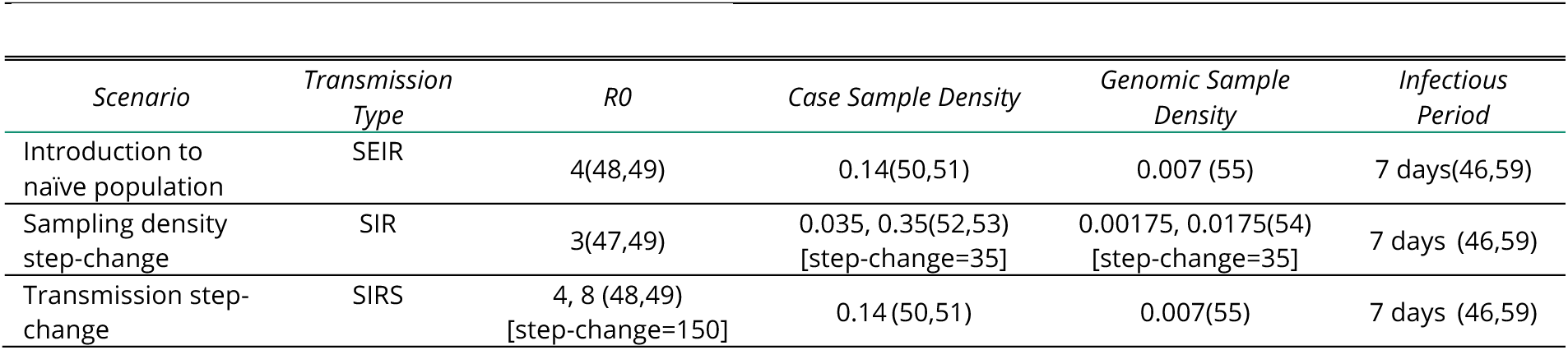
Characteristics of simulated outbreak scenarios for model validation. The numbers in parentheses indicate the literature used to support the parameter choice. ‘step-change’ refers to the time of the step-change of the parameter from the beginning of the simulation in days.

### Model Distribution and Data Availability

The model source code, example parameter files and guidance on usage are available at the GitHub repository https://github.com/ciarajudge/EpiFusion under the GNU General Public License. Code used for the specific models and plots in this manuscript is also provided. EpiFusion is also downloadable as an executable JAR file for convenient use. The program takes an XML file as input, which contains the data and parameters for the model. The user does not need to define the compartmental model (i.e. SIR, SEIR etc), but parameterisation of rates *β*, *γ*, *φ* and *ψ* is necessary with a selection of options available to users for priors or step-changes at specific times.

## 3. Results

### Comparison of EpiFusion Approaches

For the first stage of model validation, we evaluated how well EpiFusion could reconstruct trajectories of infections and Rt corresponding to simulated outbreaks that were estimated under several different scenarios. We compared the performance of EpiFusion using solely case incidence data, solely a phylogenetic tree, and using both datasets combined. The metrics by which models are compared and their statistics are summarised in Table 2.

**Table 2.** Statistics used to evaluate model performance under scenarios 1, 2, and 3 for analyses using case incidence only (epi), phylogenetic tree only (phylo), and both data sources combined (combo). The best or joint-best result for each statistic for each scenario is highlighted in bold. Trajectory RMSE: root-mean-squared error. Trajectory Coverage: proportion of true trajectory that falls within the 95% HPD, scaled by 0.95. Scaled HPD Width: mean width of the 95% highest posterior density interval, scaled by the true value. Continuous Ranked Probability Score: mean CRPS across the trajectory time series. Brier Score: Classification accuracy for transmission phase (Rt) being above or below 1. Further details on the calculation of these statistics are included in the Supplementary Information (page 43).

| Evaluation Metric | Introduction Scenario |  |  | Sampling Step-Change |  |  | Transmission Step-Change |  |  |
| --- | --- | --- | --- | --- | --- | --- | --- | --- | --- |
|  | Epi | Phylo | Combo | Epi | Phylo | Combo | Epi | Phylo | Combo |
| Infection Trajectory RMSE | **215** | 380 | 220 | **157** | 285 | 167 | 121 | 212 | **103** |
| Infection Trajectory Coverage | **0.81** | 0.62 | 0.73 | **0.99** | 0.87 | 1.02 | **0.98** | **0.98** | **1.02** |
| Scaled Infection Trajectory HPD Width | **1.28** | 21.79 | 1.39 | 1.48 | **1.28** | **1.28** | **0.35** | 0.63 | 0.39 |
| Rt Trajectory RMSE | 0.48 | 0.56 | **0.32** | 0.81 | **0.78** | 0.83 | 0.25 | **0.18** | 0.19 |
| Rt Trajectory Coverage | **1.0** | 1.02 | 1.02 | **0.86** | 0.77 | 0.75 | **0.98** | **1.02** | **0.98** |
| Scaled Rt Trajectory HPD Width | 1.25 | 2.22 | **1.06** | 1.60 | **1.47** | 1.51 | 1.10 | 0.75 | **0.61** |
| Continuous Ranked Probability Score | **88.2** | 235 | 101 | 71.2 | 115 | **64.9** | 56.3 | 117 | **51.1** |
| Rt Continuous Ranked Probability Score | 0.25 | 0.31 | **0.23** | 0.29 | 0.42 | **0.28** | 0.13 | 0.10 | **0.09** |
| Brier Score – Transmission Phase | **0.02** | 0.07 | 0.03 | **0.02** | 0.04 | **0.02** | **0.16** | 0.24 | **0.16** |

We first considered a scenario in which a novel pathogen enters an immune naïve population with constant sampling: the ‘introduction scenario’. Here, each approach successfully inferred a trajectory that was close to the true epidemic trajectory (*Figure 3*), however the tree only approach underperformed according to the metrics we chose for evaluation (*Table 2*). The case incidence only approach was most successful in fully capturing the true infection trajectory (Infection Trajectory Coverage: *81%*) compared to tree-only and combined models (*62%, 73%*) (see Table 2 and the Supplementary Information page 43 for a description of the statistics). The Continuous Ranked Probability Score (CRPS) was used to evaluate the probability of the true infection or Rt trajectory given the posterior infection or Rt trajectories from each model, where a lower value equates to a more accurate result. Here the case incidence only approach also performed best (*88.2* vs *235* and *101* for tree only and combined approaches, respectively). The reduced trajectory coverage (*73%*) in the combined approach compared to the case-incidence only approach (*81%*) resulted from a slightly out-of-phase epidemic peak, but the effect of this is diminished in the Rt estimates (*Figure 4*). Here, the approaches perform more equitably in terms of coverage, with the tree only and combined model over-covering by a very small margin (*1.02*) and the case incidence only model achieving perfect coverage (*1.0*). However, the Rt CRPS for the combined approach showed a small improvement compared to the individual approaches (*0.23* vs *0.25* and *0.31*), and also demonstrated less root-mean-square error (RMSE) (*0.32* vs *0.48* and *0.56*). These improvements in Rt estimation are accompanied by a reduction in the width of the HPD intervals (*1.06* vs *1.25* and *2.22*), a positive result indicating increased confidence provided that coverage is maintained, which it is in this case. We note that the improvement of the CRPS in the combined approach compared to the case incidence approach is marginal (*0.02*). We also used the Brier score (mean squared error between the probabilistic prediction and the true outcome) to evaluate each approach based on its ability to predict transmission phase, i.e. correctly estimating if Rt is above or below 1, where a lower Brier score indicates improved performance. We find each approach to be adept at classifying Rt as above or below 1, with the case incidence approach (*0.02*) marginally outperforming the combined approach (*0.03*), followed by the tree-only approach (*0.07*).

**Figure 3.**
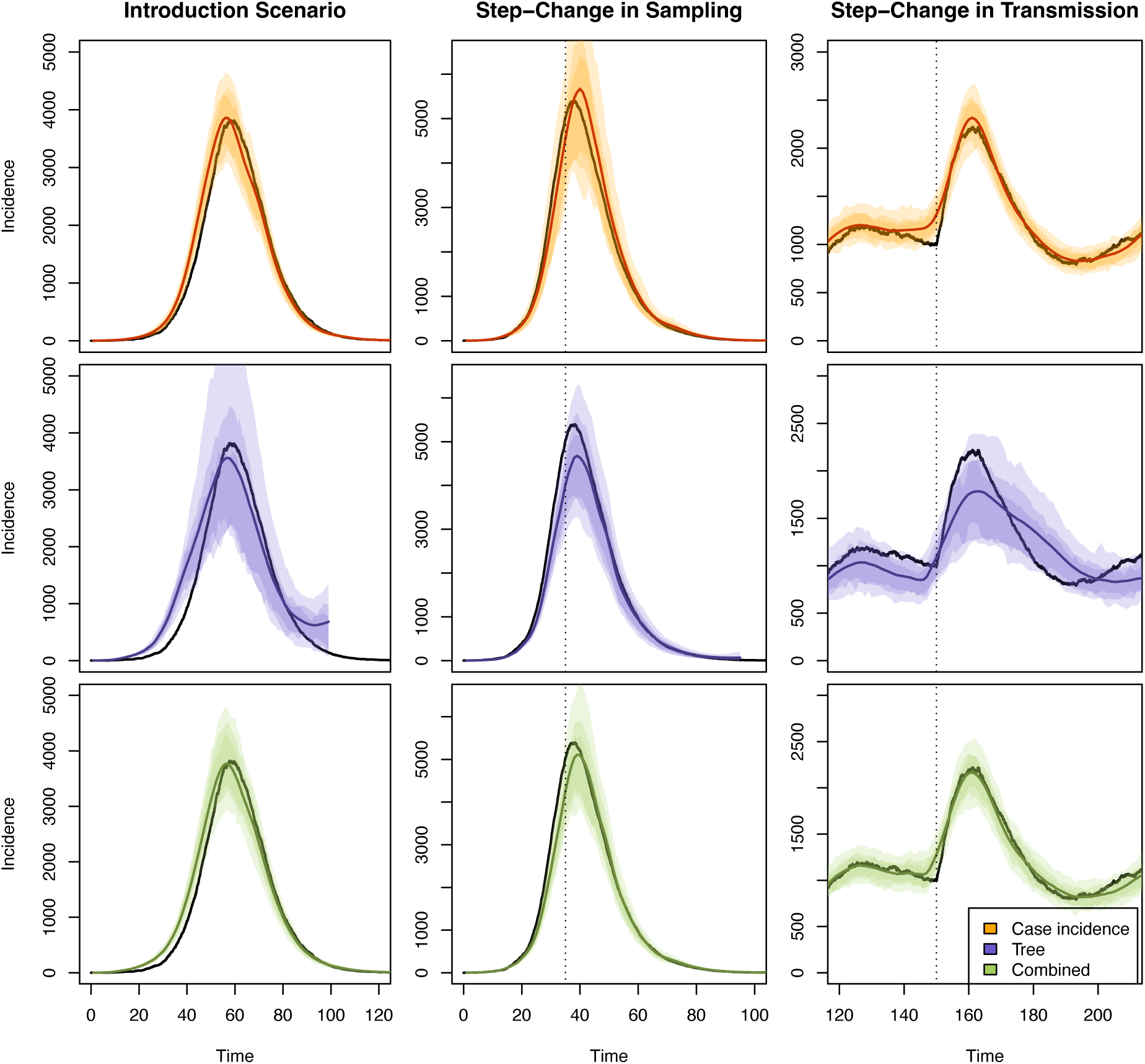
Inferred trajectories from EpiFusion using only case incidence (orange), only the phylogenetic tree (purple) and both data types combined (green) for the three scenarios tested. The true number infected over time is represented by the black line. 95%, 80% and 66% confidence intervals are represented by increasingly dark shaded regions. Times of step-changes are marked by the vertical dotted lines for the step-change in sampling and transmission scenarios.

**Figure 4.**
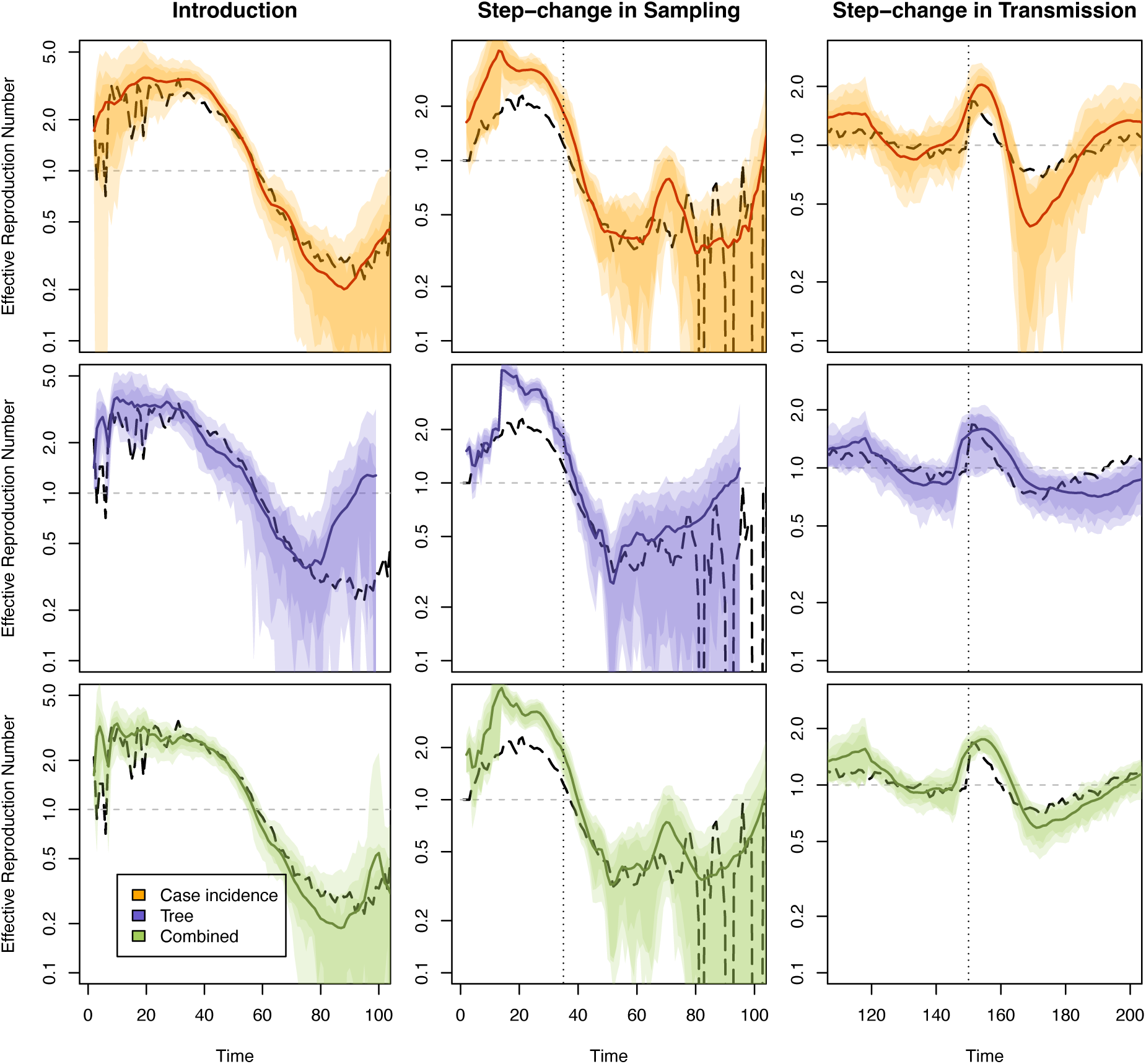
Inferred Rt from EpiFusion using only case incidence (orange), only the phylogenetic tree (purple) and both data types combined (green) for the three scenarios tested. True Rt is represented by the dashed black line. 95%, 80% and 66% confidence intervals are represented by increasingly dark shaded regions. Times of step-changes are marked by vertical dotted lines. An Rt of 1 is marked by the dashed horizontal line. The true Rt fluctuates at the end of the sampling step-change scenario due to very low prevalence as the outbreak ends.

The second scenario consisted of a simulated outbreak with similar transmission dynamics to the introduction scenario but for which levels of both genomic and case sampling are low during the initial period of spread until more widespread surveillance is introduced (thus a step-change in sampling density). Here, the date of this step-change is provided as a fixed parameter to the model under the assumption that it would be known to health authorities, but fixing this parameter is not strictly necessary to run the model. The sampling rates before and after the step-change are inferred as parameters of the MCMC.

For this analysis, all three approaches suffer from overestimating the Rt trajectory early in the simulation. We attribute this to difficulties in parameterising the sampling rates when prevalence is very low at the beginning of the simulated outbreak, and due to the dramatic sampling step-change. The case incidence approach demonstrates the best Rt trajectory coverage (*0.86*), likely due to its wider HPD intervals (*1.60*). However, the combined approach demonstrates good coverage of the true infection trajectory (*1.02*), while at the same time reducing the HPD interval width (*1.28 vs 1.48*) in comparison to the case incidence only approach (*Figure 3*). The combined approach also led to the best infection and Rt trajectory CRPS results (*64.9* and *0.28*), and demonstrated improved accuracy in predicting transmission phase compared to the tree only approach (Brier score *0.02 vs 0.04*). The sampling step-change scenario is the only scenario where the combined approach did not lead to an improvement on one or both individual approaches for all metrics, underperforming on the Rt Trajectory RMSE and coverage, however the differences between all three approaches for these metrics are marginal.

The final scenario examined was a scenario in which a step-change in transmission was simulated, where a pathogen experiencing endemic transmission undergoes a spontaneous increase in transmission, but where sampling parameters are constant. For this analysis, the date of the transmission increase was inferred as a parameter of the MCMC (it is possible to fit any number of rate step-changes with EpiFusion; it is not currently possible to infer the number of step changes). All three models broadly captured the epidemic trajectories (*Figure 3*), with equal coverage (*0.98, 0.98, 1.02*), however the combined approach resulted in the lowest trajectory RMSE (*103* vs *121* and *212*) and CRPS (*51.1* vs *56.3* and *117*). When we examine the Rt estimates the combined approach also resulted in a slightly improved CRPS here (*0.09* vs *0.13* and *0.10*), along with improved transmission phase classification (*0.16*) compared to the tree only approach (*0.24*). The approaches were then evaluated for their ability to reconstruct selected parameters that were used to simulated different scenarios (*Figure 5*).

**Figure 5.**
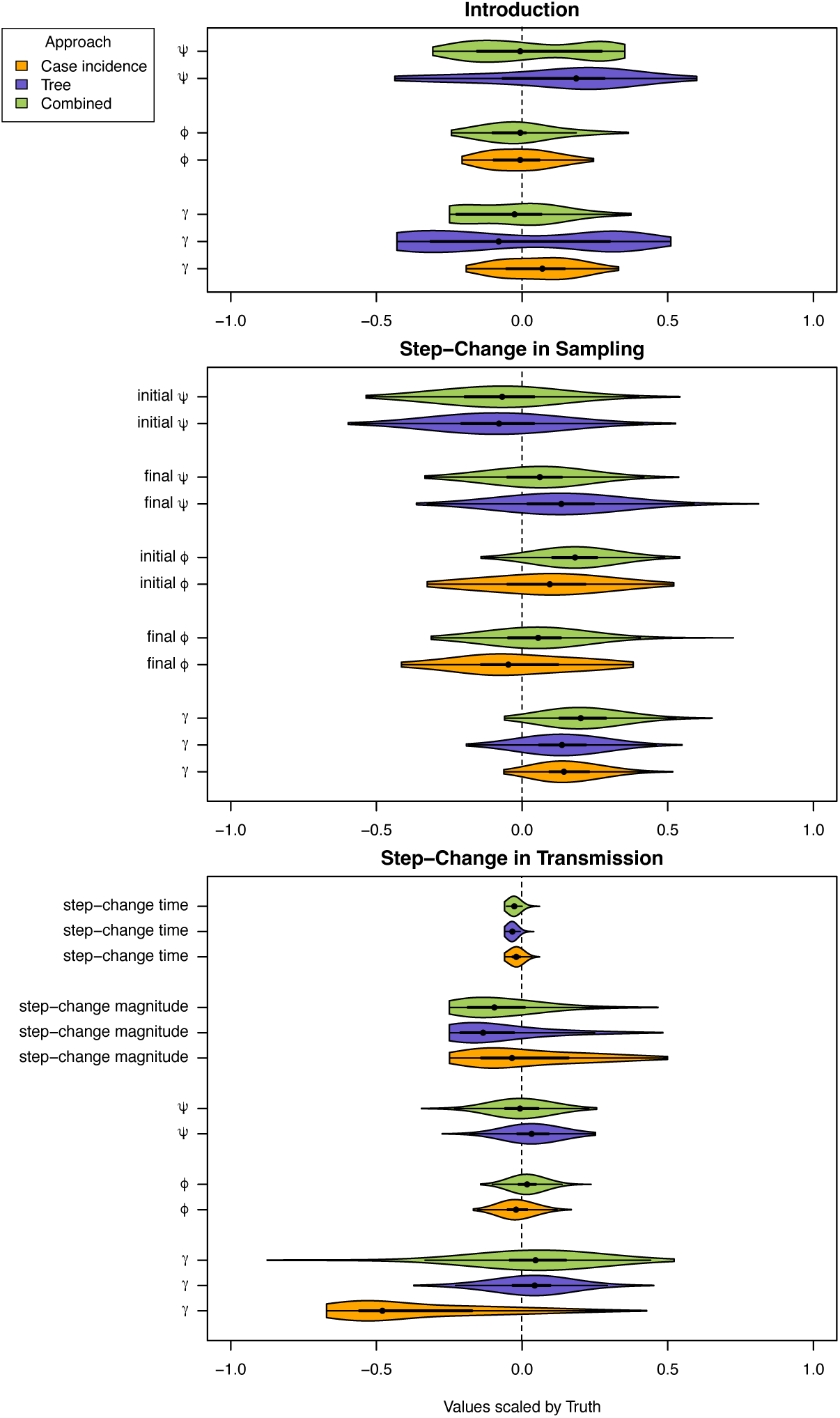
Violin plots of parameter posterior densities of the EpiFusion models, scaled by the true simulated value of the parameter and truncated at the 95% HPD boundaries. The median parameter values are marked by black circles, and the heavy black lines denote the interquartile range. The ‘true value’ is marked by the dotted black line.

While all approaches capture the true values of all parameters within the 95% HPD intervals for all scenarios, the approaches vary in bias and uncertainty. In general, all approaches agree on the direction of the bias, with the exception of *γ* in the introduction and step-change in transmission scenario. A key test for the step-change in transmission scenario is the ability of the models to detect the increase in Rt close to the true time of the change, and most importantly without any delay (*Figure 5*, ‘step-change time’). The tree-only approach predicted the uptick in transmission earliest, however it never confidently indicated the increase of Rt to greater than 1.0 (i.e. the lower HPD bound is never above 1.0), with the combined approach being the first approach to do this.

### Benchmarking against existing approaches

The second stage of model validation was to compare the performance of the EpiFusion combined data model against existing Rt inference methods (*Figure 6*). We used (i) EpiNow2 (56), (ii) a Birth-Death Skyline Serially Sampled model implemented in BEAST2 (BDSky) (14), and (iii) TimTam (30) implemented in BEAST2 to represent existing epidemiological only, phylodynamic only, and combined approaches, respectively. Table 3 provides a qualitative comparison of the capabilities and limitations of EpiFusion relative to these tools, and other models referenced in the Introduction. Further information on model specifications for these analyses are included in the Supplementary Information (page 42).

**Figure 6.**
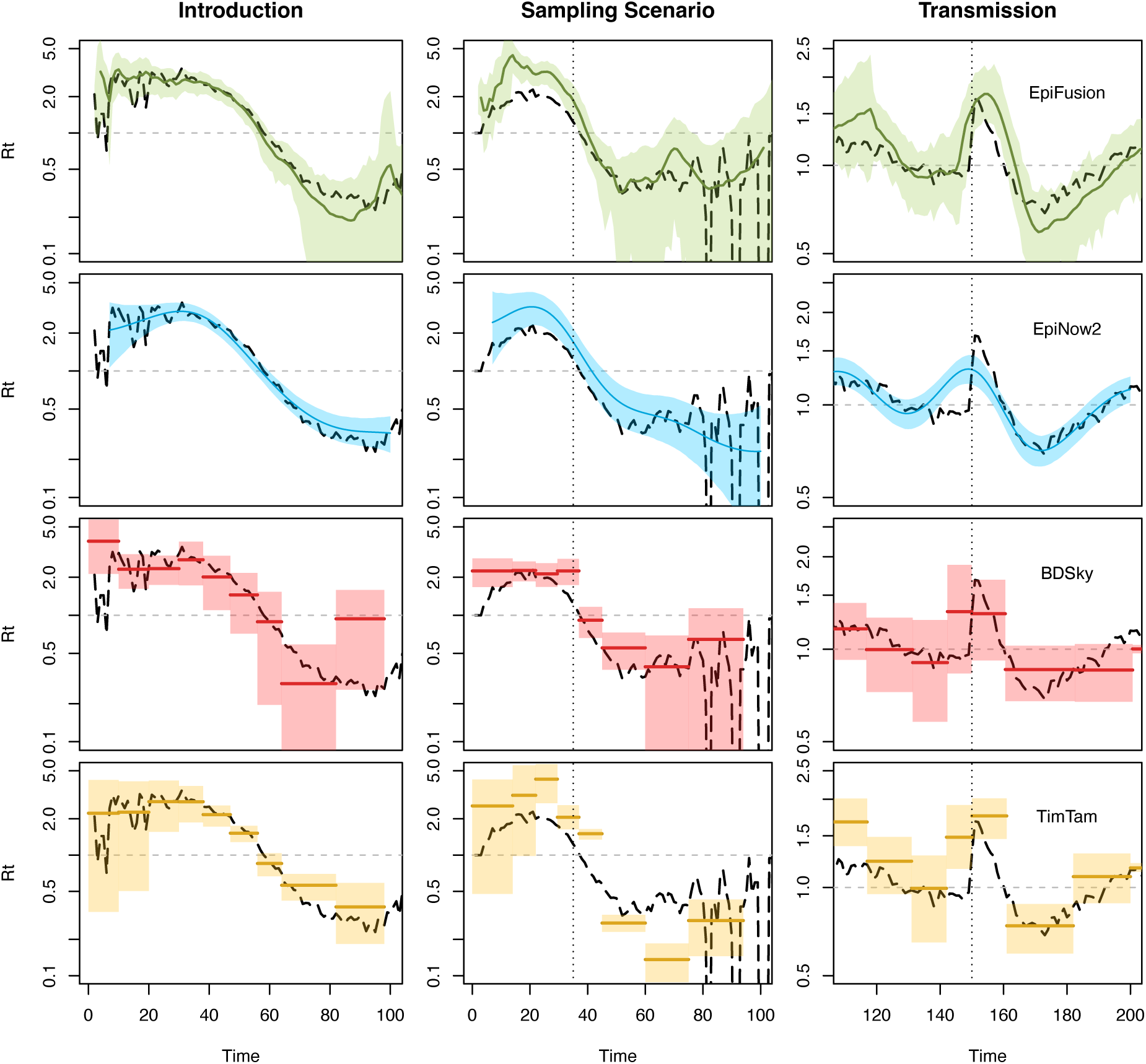
Estimated Rt for the three validation scenarios from EpiFusion (green), EpiNow2 (blue), BDSky (red) and TimTam (yellow).

**Table 3.** Comparison of the key characteristics of EpiFusion compared to the tools and literature referred to in this manuscript. Rasmussen’11 denotes Rasmussen et. al (2011), which was referenced in the introduction. However, the model is not distributed for use as a software or program, so we were unable to assess its computational efficiency (*). (BD – birth death).

|  | Uses Case Incidence | Uses Phylogenetic Tree | Infers Phylogenetic Trees | Infers Continuous Trajectories | Scalable to Large Datasets | BD Theory |
| --- | --- | --- | --- | --- | --- | --- |
| EpiFusion | ✓ | ✓ |  | ✓ | ✓ | ✓ |
| EpiNow2 | ✓ |  |  | ✓ | ✓ |  |
| EpiInf | ✓ | ✓ | ✓ | ✓ |  | ✓ |
| TimTam | ✓ | ✓ | ✓ |  | ✓ | ✓ |
| BDSky |  | ✓ | ✓ |  | ✓ | ✓ |
| Rasmussen '11 | ✓ | ✓ |  | ✓ | * |  |

Rt posteriors were obtained from each benchmarking model for all three scenarios and compared to the combined EpiFusion approach for each. The strengths and weaknesses of the different models are apparent when examining their performance under selected scoring criteria (*Table 4*).

**Table 4.** Model Benchmarking.

|  | Introduction Scenario |  |  |  | Sampling Step-Change |  |  |  | Transmission Step-Change |  |  |  |
| --- | --- | --- | --- | --- | --- | --- | --- | --- | --- | --- | --- | --- |
|  | *EpiFusion* | *EpiNow2* | *BDSky* | *TimTam* | *EpiFusion* | *EpiNow2* | *BDSky* | *TimTam* | *EpiFusion* | *EpiNow2* | *BDSky* | *TimTam* |
| Rt RMSE | 0.31 | **0.27** | 0.76 | 0.38 | 0.76 | 0.56 | **0.43** | 0.88 | 0.19 | **0.12** | 0.14 | 0.20 |
| Rt CRPS | 0.14 | **0.12** | 0.36 | 0.19 | 0.27 | 0.29 | **0.23** | 0.51 | 0.09 | **0.06** | 0.08 | 0.17 |
| Brier Score | **0.01** | **0.01** | 0.07 | 0.05 | **0.02** | 0.03 | 0.04 | 0.10 | 0.16 | **0.15** | 0.22 | 0.32 |
| Rt Coverage | **0.99** | 0.94 | 0.88 | 0.87 | **0.76** | 0.68 | 0.73 | 0.36 | **0.98** | 0.91 | 0.96 | 0.19 |

Each model captured the general trend of transmission for all three scenarios, with some weaknesses. Using EpiFusion resulted in improved calibrated coverage for all three scenarios. EpiFusion also led to better Brier scores compared to other methods for the introduction and sampling scenarios, however the difference between EpiNow2 and EpiFusion for this metric was consistently marginal. EpiFusion did not lead to overall improvements for Rt RMSE or CRPS, but notably never produced the worst performance under any scenario and metric combination. The EpiNow2 model inferred relatively accurate Rt trajectories with high certainty (*Figure 6*) and performed well in most scenarios under all scoring criteria, particularly the introduction and transmission step-change scenarios. The model somewhat struggled with identifying sharp fluctuations in transmission, possibly due to the smoothing influence of the gaussian process. The BDSky approach particularly underperformed in the introduction scenario according to our criteria but resulted in improved RMSE and CRPS compared to all other approaches in the sampling step-change scenario. Conversely, TimTam struggled with the sampling scenario and transmission scenario, particularly in the coverage scores due to the limitations of its narrow HPD intervals and piecewise constant estimates. However, Figure 6 shows that the model quite clearly detects a change of transmission (signalled by a jump in the inferred Rt and posteriors), despite not accurately inferring the actual values of Rt.

## 4. Discussion

We outline EpiFusion, a computationally tractable and flexible infrastructure for the combination of phylogenetic and epidemiological data to estimate infection and Rt trajectories. EpiFusion fills a gap in current modelling approaches at the intersection between the fields of phylodynamics and epidemiology (*Table 3*). We show that by combining data types with EpiFusion it is often possible to improve the accuracy of Rt or infection trajectory estimates compared to one or both individual approaches.

The introduction scenario aimed to represent a situation such as the emergence of a novel pathogen (60) or the expansion of an existing pathogen into a new ecological niche (61). All three EpiFusion approaches (case incidence only, tree only, combined) were able to accurately reconstruct the epidemic trajectories of the simple, single epidemic peak, with the case incidence only approach resulting in the greatest trajectory coverage. For all three approaches the trajectory is shifted earlier with roughly the same bias, this may be due to difficulties in modelling SEIR transmission (particularly for the tree only and combined approaches). For this scenario, with smooth changes in transmission and constant sampling, the performance was broadly comparable between the three approaches, however the tree only approach underperformed slightly. Furthermore, the inference using the phylogenetic tree is truncated at day 98 of the simulation as this is the date of the last sampling event on the tree. An advantage of the combined model is that the trajectories can be jointly inferred up until this time, but the estimates can continue until the present time using the case incidence data only.

We subsequently considered more complex scenarios of changes in sampling and transmission rate, which better reflect the reality of infectious disease outbreaks but which are widely acknowledge to complicate the estimation of Rt. This allowed us to examine how combining phylodynamic and epidemiological models and data could improve our ability to accurately estimate Rt under such challenging scenarios. The rationale for the step-change in sampling scenario was to emulate a pathogen that begins circulation initially undetected by health authorities, resulting in a lack of data from the early stages of an outbreak and a lack of comparability in case numbers before and after detection is scaled up. This also applies for novel pathogens that do not have established means of clinical diagnoses or reporting, or where testing is initially limited. For example, during the Zika virus epidemic in Brazil in 2016, case detection rates rose sharply following the implementation of widespread PCR testing (53), compared to the beginning of the outbreak. Each of the models struggled with overestimation of Rt before the sampling step-change – we note that in the simulated data, the change in sampling density was instant, whereas a sharp but not completely instantaneous scale up might be more realistic and easier to capture in the models. The tree-only approach demonstrated more advantages during this scenario than in the other scenarios tested, which is likely due to the additional information captured by birth events in the tree even when sampling was low. Notably the combined approach led to improved infection trajectory and Rt continuous ranked probability scores, the probabilistic scoring rule we chose for model comparison. The combined model also outperformed the individual approaches according to the Brier score metric. This indicates that the combined approach may benefit estimation of whether an epidemic is growing or declining, which is a useful public health tool to be able to evaluate with certainty (62,63). By this metric the combined model also outperformed the individual approaches.

The step change in transmission scenario was used to mimic a sudden increase in transmission, such as a change in human behaviour (e.g. school holidays end, non-pharmaceutical intervention ceases), or a change in the intrinsic transmissibility of a pathogen (e.g. a new variant (64)). The phylogenetic tree simulated from ReMASTER is more applicable to the former, in that all ‘active’ lineages at the time of the step-change undergo an equal increase in transmission which is not what would be observed in the case of a new, more transmissible variant. Currently, EpiFusion does not attempt to infer lineage specific transmission rates, so this distinction is not important, but any future incorporation of lineage specific analyses will require this to be considered. Notably, the tree-only approach was the earliest approach to note the simulated uptick in transmission (*Figure 4*), but the lower bound of the Rt HPDs did not increase above 1 for the approach at any point. Conversely, the case incidence only approach is slightly delayed in confidently indicating an increase of Rt above 1 compared to the other approaches, does capture the magnitude of the transmission increase. The combined approach achieves a good balance, confidently inferring time and magnitude of the increase of transmission, particularly in the inferred infection trajectories (*Figure 3*). This further demonstrates the value of combining data types: uncertainty from the weekly resolution of the case reporting propagates through to the epidemiological model predictions, whereas branching events in the phylogenetic tree can capture early changes in transmission dynamics. This is also visible in examining the performance metrics in Table 2, where the combined approach led to the best result for seven out of nine metrics. Again, the combined model led to improved continuous ranked probability scores for both the infection and Rt trajectories, and an improved Brier score compared to the tree only analysis.

Overall, the combined-model tended to reduce uncertainty compared to case-only and phylogenetic-only approaches, as observed by narrowing of the HPD intervals of the infection trajectories, while maintaining coverage (*Table 2*). For all three scenarios, the combined approach led to the best Rt CRPS and frequently outperformed the individual approaches according to our other metrics. There are some circumstances, however, where either the pure epidemiological or phylodynamic approaches are preferable, such in the introduction scenario, where the case incidence only approach performed best. This points to the benefit of the versatility of the EpiFusion program; while we emphasise the combined inference abilities of EpiFusion, it is possible to run analyses using either case incidence or the phylogenetic tree alone. Furthermore, the program is sufficiently fast for users to test tree-only, incidence only, and combined approaches in a reasonable timeframe and subsequently choose an approach according to their specific requirements. It is also theoretically possible to specify the weight of each dataset’s contribution to the inference, allowing further customisation of the combined approach. Going forward, we aim to characterise the implementation and effect of data weighting more thoroughly.

Our benchmarking of EpiFusion’s combined model against existing approaches shows that the model can achieve comparable or improved results compared to established epidemiological or phylodynamic tools for these three scenarios, however how this generalises to different situations and real data must be further examined. For many performance metrics used the difference in scores between all models was marginal, but we did not necessarily expect major improvements with EpiFusion as the model is still in early development. However, EpiFusion led to improved Rt trajectory coverage in all scenarios compared all other models (*Table 4).* It should be noted that the BDSky and TimTam models take a sequence alignment as their input, and infer tree structure throughout the inference process, whereas EpiFusion does not currently incorporate phylogenetic uncertainty or infer trees and took the true ReMASTER simulated tree as its input; thus conferring a marked advantage. This ‘true tree’ will not be available for real-world data, and a time-scaled phylogenetic tree will need to be assembled from genomic sequence data, thus introducing phylogenetic uncertainty that will likely impact the performance of EpiFusion. The impact of this uncertainty, and how best to accommodate it in EpiFusion, will be further characterised in future publications. EpiNow2 proved difficult to parameterise, particularly for the sampling and transmission scenarios, so it is also possible that improved parameterisation of the model would result in better estimates.

Our approach currently has limitations and necessitates some assumptions that provide opportunity for future improvements. Our evaluation of EpiFusion performance thus far is limited to simulated datasets, so an essential next step will be a case study on a real world dataset. We will use this case study to examine how this model generalises to real outbreak data, and further characterise the best use-case for this model.

Unlike other recent approaches, EpiFusion does not estimate phylogenies alongside trajectories, and instead takes phylogenetic trees as inputs. This is a limitation that we aim to address by working to incorporate phylogenetic uncertainty in the future. However, the computational trade-off of not performing tree inference means our method may be appropriate for use in rapidly unfolding outbreaks once it has been further validated in a real-time setting, as it is highly scalable to inclusion of trees with thousands of tips.

The lightweight composition of this model provides the opportunity for the introduction of additional complexity without overtly increasing computational load. This includes the introduction of population structure or vector population dynamics. The separation of the phylogenetic and epidemiological observation models in EpiFusion also lays the foundation for the combination mathematical epidemiological models that previously would have been too complex to integrate into the phylodynamic likelihood with phylogenetic data to jointly model epidemic trajectories.

In conclusion, we propose EpiFusion as a new addition to the small but growing number of tools that integrate phylodynamics and epidemiology for the modelling of infectious disease. EpiFusion builds upon the foundation laid by its predecessors to make improvements in computational efficiency, temporal resolution and flexibility. The intended future development for this work is to demonstrate this approach on real-world data, making the necessary steps forward in complexity to accommodate this while investigating potential improvements such as data-weighting or population structure.

## Acknowledgements

The authors would like to express their gratitude to Dr. Alex Zarebski, Prof. Oliver Pybus, Prof. Katia Koelle, Dr. David Hodgson, Dr. Alexis Robert, Emilie Finch and Ciara McCarthy for their advice and guidance during the development of this work.

## Funding Statement

CJ was supported by a Bloomsbury Colleges PhD Studentship and a National University of Ireland Denis Phelan Scholarship. TWR was supported by funding from the Wellcome Trust (grant: 20650/Z/17/Z). SA was funded by the Wellcome Trust (grant: 210758/Z/18/Z). TS received funding from the European Research Council (ERC) under the European Union’s Horizon 2020 research and innovation programme grant agreement no. 101001077. Further, TS acknowledges funding from ETH Zurich. OJB was supported by a UK Medical Research Council Career Development Award (MR/V031112/1). SCH was supported by a Sir Henry Wellcome Postdoctoral Fellowship from the Wellcome Trust (220414/Z/20/Z) [https://welcome.org/]. For the purpose of open access, the authors have applied a CC BY public copyright licence to any Author Accepted Manuscript version arising from this submission.

## Supplementary Information

### Importance Sampling Implementation

To prevent the simulation of trajectories which are incompatible with the phylogenetic tree we implement importance sampling. Importance sampling is a statistical technique used to efficiently estimate properties of a model, by focusing on more relevant samples. This results in a subtle change to the exact implementation of the process model (*Main Text Eqn 1*) where the terms are adjusted based on the structure of the phylogenetic tree. Specifically, the infection rate, *β*_t_, is first divided into *observed* and *unobserved* infection rates *β°* and *β^u^ (Eqns 6&7).* The rate *γ* is adjusted to prevent recovery events that violate the phylogenetic tree structure (i.e., that lead to estimates of the number of infections that are lower than the number of concurrent viral phylogenetic lineages, *l*), resulting in *allowed* and *forbidden* recovery rates *γ*^-^ and *γ*\* *(Eqns 8&9)*.

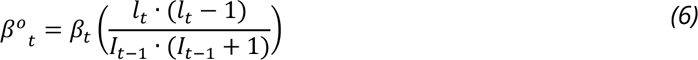

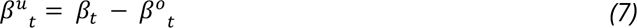

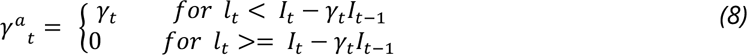

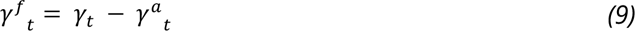

These equations give the model for epidemic trajectory simulation under importance sampling:

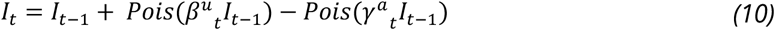

### ReMASTER Data Simulation Parameters

**Table S1.** Introduction Scenario Simulation Parameters.

| Table S1 – Introduction Scenario Simulation Parameters |  |  |  |
| --- | --- | --- | --- |
| Initial Populations |  | Rates |  |
| S | 19999 | I + S -> I + E | 0.000029 |
| E | 0 | E -> I | 0.2 |
| I | 1 | I -> R | 0.143 |
| R | 0 | I -> sample | 0.02 |
| sample | 0 |  |  |

**Table S2.** Sampling Scenario Simulation Parameters.

| Table S2 – Sampling Scenario Simulation Parameters |  |  |  |
| --- | --- | --- | --- |
| Initial Populations |  | Rates |  |
| S | 19999 | I + S -> I + I | 0.00002 |
| I | 1 | I -> R | 0.143 |
| R | 0 | I -> sample (T < 35) | 0.005 |
| sample | 0 | I -> sample (T >= 35) | 0.05 |

**Table S3.**
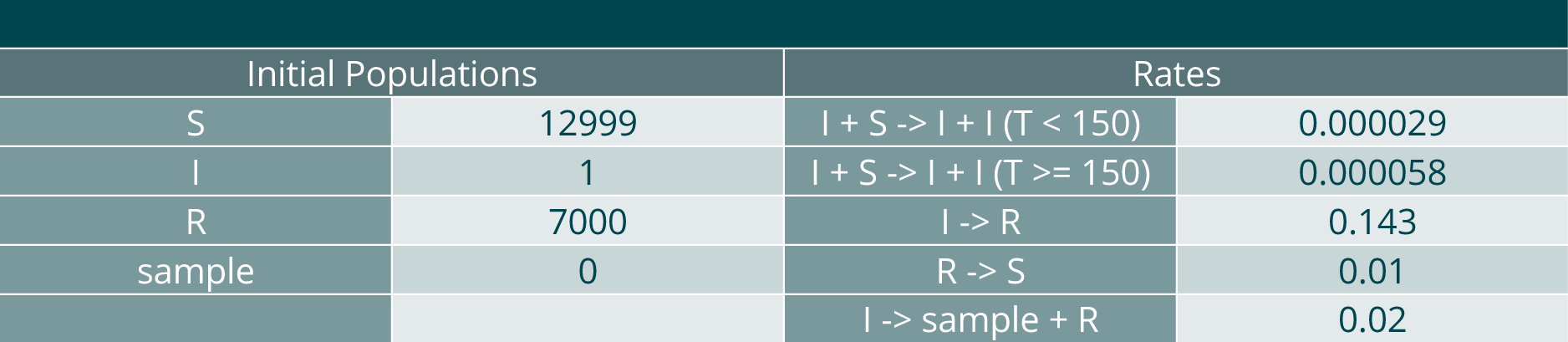
Transmission Scenario Simulation Parameters.

| Table S3 – Transmission Scenario Simulation Parameters |  |  |  |
| --- | --- | --- | --- |
| Initial Populations |  | Rates |  |
| S | 12999 | I + S -> I + I (T < 150) | 0.000029 |
| I | 1 | I + S -> I + I (T >= 150) | 0.000058 |
| R | 7000 | I -> R | 0.143 |
| sample | 0 | R -> S | 0.01 |
|  |  | I -> sample + R | 0.02 |

### Simulated Data – Scenarios 2 and 3

**Supplementary Figure S1.**
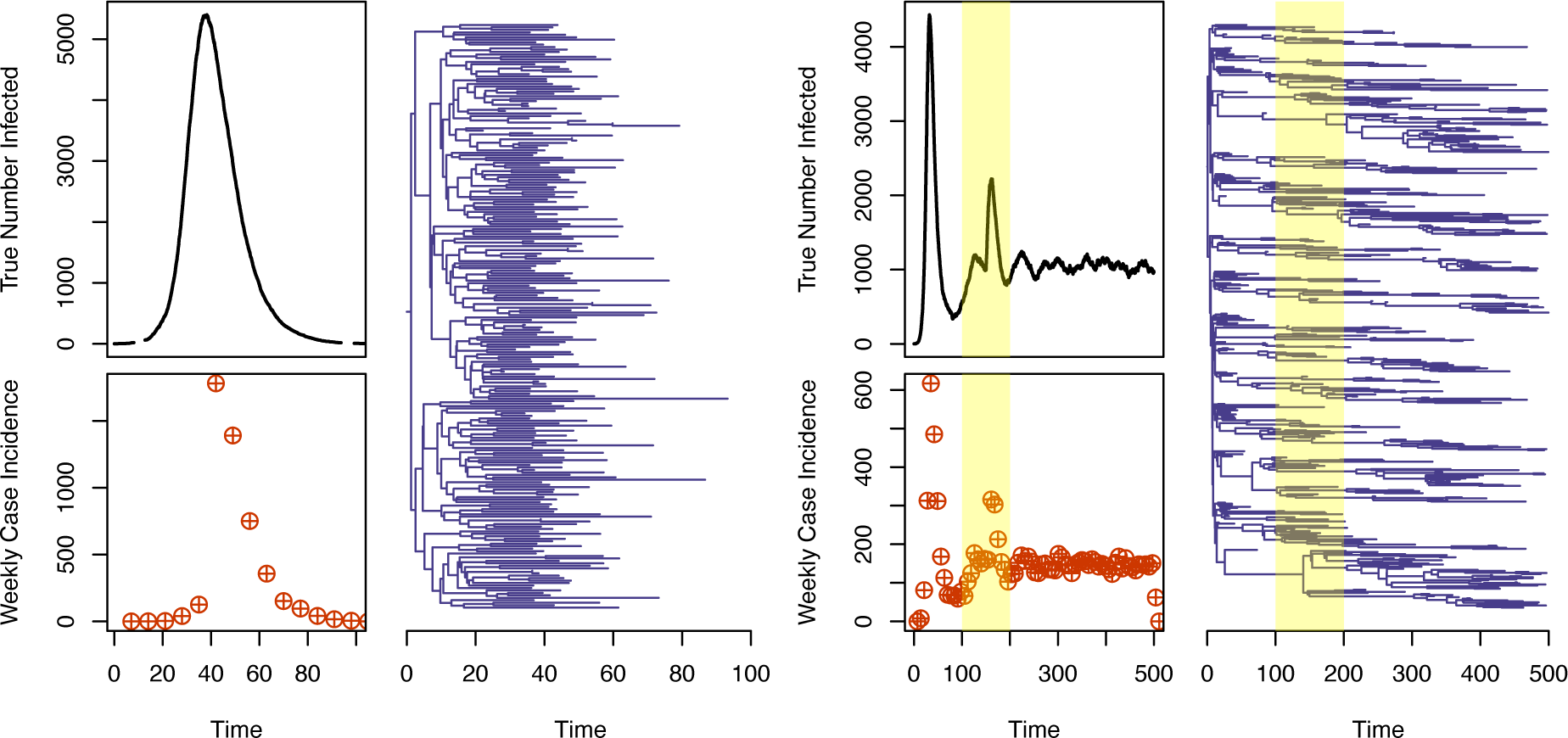
Simulated datasets for the step-change in sampling (left) and transmission (right) scenarios. True infection trajectory is shown as a black line, with weekly case incidence shown as orange points. The phylogenetic tree is shown in purple. Note, for the transmission scenario (right) we focus on day 100-200 of this simulation for this paper (highlighted in yellow). ReMASTER simulations require initiation with only one infected person, but we wished to examine a scenario with endemic transmission, so we initiate the simulation and introduce the step change after equilibrium has been reached at ∼day 100).

**Table S4.** Model Priors Validation Stage 1.

|  | Approach | Priors |  |  |  |  |  |  |  |
| --- | --- | --- | --- | --- | --- | --- | --- | --- | --- |
| | | $\gamma$ | $\psi$ | $\varphi$ | $\beta_1$ | $\beta_\sigma$ | $T_{sc}$ | $M_{sc}$ | $I_{init}$ |
| Introduction | Case Incidence | Truncated Norm<br>Mean = 0.14<br>StdDev = 0.04<br>Lowerbound = 0 | NA | Truncated Norm<br>Mean = 0.14<br>StdDev = 0.03<br>Lowerbound = 0 | Uniform<br>0.2-0.6 | Truncated Norm<br>Mean = 0.02<br>StdDev = 0.02<br>Lowerbound = 0 | NA | NA | NA |
|  | Tree | Truncated Norm<br>Mean = 0.14<br>StdDev = 0.04<br>Lowerbound = 0 | Truncated Norm<br>Mean = 0.001<br>StdDev = 0.0003<br>Lowerbound = 0 | NA | Uniform<br>0.2-0.6 | Truncated Norm<br>Mean = 0.02<br>StdDev = 0.02<br>Lowerbound = 0 | NA | NA | NA |
|  | Combined | Truncated Norm<br>Mean = 0.14<br>StdDev = 0.04<br>Lowerbound = 0 | Truncated Norm<br>Mean = 0.001<br>StdDev = 0.0003<br>Lowerbound = 0 | Truncated Norm<br>Mean = 0.14<br>StdDev = 0.03<br>Lowerbound = 0 | Uniform<br>0.2-0.6 | Truncated Norm<br>Mean = 0.02<br>StdDev = 0.02<br>Lowerbound = 0 | NA | NA | NA |
| Sampling Step-Change | Case Incidence | Truncated Norm<br>Mean = 0.143<br>StdDev = 0.03<br>Lowerbound = 0 | NA | Truncated Norm<br>Mean = 0.005, 0.05<br>StdDev = 0.001, 0.01<br>Changetime = 35 | Uniform<br>0.1-0.8 | Truncated Norm<br>Mean = 0.05<br>StdDev = 0.01<br>Lowerbound = 0 | NA | NA | NA |
|  | Tree | Truncated Norm<br>Mean = 0.143<br>StdDev = 0.03<br>Lowerbound = 0 | Truncated Norm<br>Mean = 0.00025, 0.0025<br>StdDev = 0.00005, 0.0005<br>Changetime = 35 | NA | Uniform<br>0.1-0.8 | Truncated Norm<br>Mean = 0.05<br>StdDev = 0.02<br>Lowerbound = 0 | NA | NA | NA |
|  | Combined | Truncated Norm<br>Mean = 0.143<br>StdDev = 0.03<br>Lowerbound = 0 | Truncated Norm<br>Mean = 0.00025, 0.0025<br>StdDev = 0.00005, 0.0005<br>Changetime = 35 | Truncated Norm<br>Mean = 0.005, 0.05<br>StdDev = 0.001, 0.01<br>Changetime = 35 | Uniform<br>0.1-0.8 | Truncated Norm<br>Mean = 0.05<br>StdDev = 0.01<br>Lowerbound = 0 | NA | NA | NA |
| Transmission Step-Change | Case Incidence | Truncated Norm<br>Mean = 0.143<br>StdDev = 0.02<br>Lowerbound = 0 | NA | Truncated Norm<br>Mean = 0.02<br>StdDev = 0.001<br>Lowerbound = 0 | Uniform<br>0.05-0.3 | Uniform<br>0.001-0.05 | Uniform<br>140-160 | Uniform<br>1.5x-3x | Poisson<br>Mean = 500 |
|  | Tree | Truncated Norm<br>Mean = 0.143<br>StdDev = 0.02<br>Lowerbound = 0 | Truncated Norm<br>Mean = 0.001<br>StdDev = 0.0001<br>Lowerbound = 0 | NA | Uniform<br>0.05-0.3 | Uniform<br>0.001-0.05 | Uniform<br>140-160 | Uniform<br>1.5x-3x | Poisson<br>Mean = 500 |
|  | Combined | Truncated Norm<br>Mean = 0.143<br>StdDev = 0.02<br>Lowerbound = 0 | Truncated Norm<br>Mean = 0.001<br>StdDev = 0.0001<br>Lowerbound = 0 | Truncated Norm<br>Mean = 0.02<br>StdDev = 0.001<br>Lowerbound = 0 | Uniform<br>0.05-0.3 | Uniform<br>0.001-0.05 | Uniform<br>140-160 | Uniform<br>1.5x-3x | Poisson<br>Mean = 500 |

$\gamma$ – become uninfected rate; $\psi$ – genomic sequence sampling rate (daily); $\varphi$ – case sampling rate (weekly);
$\beta_1$ – initial value of beta; $\beta_\sigma$ – standard deviation of proposal distribution for beta random walk;
$T_{sc}$ – Time of step change (days); $M_{sc}$ – Magnitude of step change; $I_{init}$ – Initial I

**Table S5.** Model Results for Validation Stage 1.

|  | Approach |  | Parameter Posteriors |  |  |  |  |  |  |  |
| --- | --- | --- | --- | --- | --- | --- | --- | --- | --- | --- |
| | | | $\gamma$ | $\psi$ | $\varphi$ | $\beta_1$ | $\beta_\sigma$ | $T_{sc}$ | $M_{sc}$ | $I_{init}$ |
| Introduction | True Values |  | 0.143 | 0.001 | 0.14 | 0.572 | NA* | NA | NA | NA |
|  | Case Incidence | Val | 0.15 (0.12, 0.18) | NA | 0.138 (0.12, 0.17) | 0.371 (0.2, 0.46) | 0.02 (0.01, 0.04) | NA | NA | NA |
| | | $\hat{R}$ | 1.016 | NA | 1.016 | 1.017 | 1.018 | NA | NA | NA |
|  | Tree | Val | 0.14 (0.08, 0.2) | 0.001 (0.0006,0.0015) | NA | 0.352 (0.2,0.486) | 0.03 (0.019, 0.04) | NA | NA | NA |
| | | $\hat{R}$ | 1.018 | 1.018 | NA | 1.018 | 1.018 | NA | NA | NA |
|  | Combined | Val | 0.14 (0.11, 0.17) | 0.001 (0.0008,0.0012) | 0.138 (0.11, 0.17) | 0.329 (0.25, 0.38) | 0.016 (0.006, 0.028) | NA | NA | NA |
| | | $\hat{R}$ | 1.017 | 1.018 | 1.016 | 1.017 | 1.018 | NA | NA | NA |
| Sampling Step-Change | True Values |  | 0.143 | 0.00025, 0.0025 | 0.005, 0.05 | 0.429 | NA* | NA | NA | NA |
|  | Case Incidence | Val | 0.17 (0.14, 0.19) | NA | 0.005 (0.004, 0.007),<br>0.49 (0.032, 0.067) | 0.52 (0.33, 0.72) | 0.043 (0.03,0.06) | NA | NA | NA |
| | | $\hat{R}$ | 1.004 | NA | 1.012, 1.011 | 1.011 | 1.01 | NA | NA | NA |
|  | Tree | Val | 0.16 (0.13, 0.19) | 0.0002(0.0001,0.0003),<br>0.0028(0.002,0.004) | NA | 0.47 (0.26, 0.67) | 0.036 (0.022,0.056) | NA | NA | NA |
| | | $\hat{R}$ | 1.0003 | 1.002, 1.019 | NA | 1.004 | 1.004 | NA | NA | NA |
|  | Combined | Val | 0.173 (0.14, 0.20) | 0.0002(0.0001,0.0003),<br>0.0026(0.002,0.003) | 0.006 (0.005, 0.007),<br>0.05 (0.038, 0.063) | 0.45 (0.23, 0.67) | 0.041 (0.025, 0.06) | NA | NA | NA |
| | | $\hat{R}$ | 1.000 | 1.004, 1.001 | 1.001, 1.001 | 1.007 | 1.003 | NA | NA | NA |
| Transmission Step-Change | True Values |  | 0.143 | 0.001 | 0.02 | NA* | NA* | 150 | 2 | 500 |
|  | Case Incidence | Val | 0.09 (0.05, 0.16) | NA | 0.02 (0.018, 0.021) | 0.13 (0.06, 0.22) | 0.015 (0.01, 0.03) | 146.86 (141,151) | 2.05 (1.5,2.79) | 494.34 (455, 535) |
| | | $\hat{R}$ | 1.017 | NA | 1.003 | 1.017 | 1.013 | 1.001 | 1.012 | 1.001 |
|  | Tree | Val | 0.148 (0.114, 0.177) | 0.001 (0.0008,0.0012) | NA | 0.178 (0.106, 0.241) | 0.011 (0.004, 0.018) | 144.87 (141, 147) | 1.82 (1.5, 2.51) | 498.23 (456, 536) |
| | | $\hat{R}$ | 1.007 | 1.004 | NA | 1.011 | 1.015 | 1.0004 | 1.009 | 1.001 |
|  | Combined | Val | 0.148 (0.110, 0.194) | 0.001 (0.0009, 0.0012) | 0.02 (0.19,0.22) | 0.165 (0.067,0.242) | 0.013 (0.01, 0.02) | 145.81 (141, 149) | 1.85 (1.51, 2.4) | 500.28 (451, 538) |
| | | $\hat{R}$ | 1.011 | 1.004 | 1.003 | 1.015 | 1.013 | 1.001 | 1.009 | 1.002 |

Means and 95% HPD Intervals in brackets of parameters inferred via MCMC (Val). Potential scale reduction factors to indicate convergence. Parameters marked by \* are used for the MCMC fitting but have no 'true
values' from the simulation process with which to compare. $\hat{R}$ : potential scale reduction factor, values close to 1 (<1.05 indicate parameter convergence)

## Model Specifications – Validation Stage 2

### EpiNow2 Parameterisation

EpiNow2 used to analyse the simulated weekly case incidence for each scenario. The reporting delay was calculated using the reporting_delay() function with a distribution according to the same parameters under which the data was simulated. The generation time was parameterised using a probability mass function of the generation time using the known values from the data simulation process, and input to the epinow function using generation_time_opts(). The epinow() function was used with default parameters, and a horizon of 0. Code used for this section is included in the project repository in the ‘publication’ folder.

### BDSky Parameterisation

Packages and software used: BDSKY 1.5.0 package (14) with BEAST 2.7 (65). XML files for these runs and log files of the results are included in the project GitHub repository in the ‘publication/benchmarking’ folder. All analyses were run using a strict clock with a prior equal to the known value from the sequence simulation, and a JC69 Site Model. Convergence was assessed using Tracer.

### TimTam Parameterisation

Packages and software used: TimTam 0.3.1 (30) package with BEAST. 2.6.7 (65). XML files for these runs and log files of the results are included in the project Github repository in the ‘publication/benchmarking’ folder. All analyses were run using a strict clock with a prior equal to the known value from the sequence simulation, and a JC69 Site Model. The analyses loosely followed the following tutorials on the TimTam Github Wiki: ‘Variable Parameters for an SIR Epidemic I’ and ‘Variable Parameters for an SIR Epidemic II’. Convergence was assessed using Tracer.

## Calculation methods for Table 2 and 3 statistics

**Table S6.** Calculation methods for Table 2 and 3.

| Measure | Method |
| --- | --- |
| RMSE | Root mean squared error between true simulated daily trajectory and the mean inferred daily trajectory from a model, calculated using the `rmse()` function from the R package `Metrics` (66). |
| Coverage | Daily upper and lower bounds of the 95% HPD interval from sampled trajectories were calculated using the `HDInterval` package in R (67), function `hdi()`. Coverage refers to the proportion over the whole trajectory that the daily true value falls within the 95% confidence interval of the inferred trajectories from the model, scaled by 0.95. For this metric, a measure closest to 1.0 indicates a better result. |
| Scaled HPD Width | HPD intervals were calculated for inferred trajectories as above. The width of the interval for each day was calculated by subtracting the upper from the lower value, and scaled by division by the true value for each day. |
| Continuous Ranked Probability Score | Continuous ranked probability score, calculated using the `crps_sample()` function of the `scoringutils` (68) package in R. The score consists of the mean daily probability of the true value given the distribution of daily values sampled in the posterior trajectories. |
| Brier Score | The $R_t$ trajectory posteriors were used to give the probability of $R_t$ being greater or less than 1 over time, this classification was compared to the truth using the `brier_score()` function of the R package `scoringutils`. |

## Notes

### Competing Interest Statement

The authors have declared no competing interest.

